# Engineering probiotic *Escherichia coli* Nissle 1917 to block transfer of multiple antibiotic resistance genes by exploiting a type I CRISPR-Cas system

**DOI:** 10.1101/2024.04.01.587504

**Authors:** Mengdie Fang, Ruiting Zhang, Chenyu Wang, Zhizhi Liu, Mingyue Fei, Biao Tang, Hua Yang, Dongchang Sun

## Abstract

Many multidrug-resistant (MDR) bacteria evolved through accumulation of antibiotic-resistance genes (ARGs). Although the potential risk of probiotics as reservoirs of ARGs has been recognized, strategies for blocking transfer of ARGs while using probiotics have rarely been explored. The probiotic *Escherichia coli* Nissle 1917 (EcN) has long been used for treating intestinal diseases. Here, we showed frequent transfer of ARGs into EcN both *in vitro* and *in vivo*, raising its potential risk of accumulating antibiotic resistance. Given that no CRISPR-Cas system is found in natural EcN, we integrated the endogenous type I-E CRISPR-Cas system derived from *E. coli* BW25113 into EcN, and showed that the engineered EcN was able to efficiently cleave multiple ARGs (i.e., *mcr-1*, *bla*_NDM-1_ and *tet*(X)). By co-incubation of EcN expressing Cas3-Cascade and that expressing Cas9, we showed that the growth of the former strain outcompeted the latter strain, demonstrating better clinical application prospect of EcN expressing the type I-E CRISPR-Cas system. Finally, the engineered EcN exhibited immunity against transfer of targeted ARGs in the intestine of a model animal (i.e. zebrafish). Our work provides a new strategy for restricting transfer of ARGs in EcN, paving the way for safe use of this probiotic and development of probiotics as living therapeutics.

## Introduction

Antimicrobial resistance (AMR) has become a global health crisis, accounting for a million annual human deaths, which is estimated to increase to 10 million by 2050^1–3^. Horizontal transfer of antibiotic resistance genes (ARGs) is the main reason for the development of AMR in bacteria^4^. Polymyxin, tetracycline and carbapenems were considered as last resort antibiotics for combating MDR bacteria. Whereas, ARGs (*mcr-1*, *bla*_NDM-1_ and *tet*X) resistant to last-resort antibiotics have been found widespread in humans and animals, as well as in the environment^5, 6^. Serving as a reservoir of ARGs, the gut microbiome could potentially horizontally transfer ARGs to pathogens and facilitate the development of MDR bacteria. Probiotics^7^, which was traditionally used for restoring microbiome balance after perturbation by antibiotics, have been considered as a substitute for antibiotics. Genetically engineered probiotics have also been used as living therapeutics for treating diseases beyond bacterial infection^8^. Despite the wide application of probiotics for a long time, their potential risk in spreading ARGs remains insufficiently investigated. Recent studies reveal that ARGs can transfer between pathogens and commensals in intestines^9^, raising a safety issue regarding potential impact of probiotics in spreading ARGs.

Traditional strategies combating AMR, such as biocide, DNA intercalating agents, and antibiotics can result in survival stress to the whole gut microbiome, which fosters development of new resistance bacteria^10^. The CRISPR-Cas system is particularly attractive in combating AMR, since it eliminates ARGs or kills only targeted bacteria by cleaving their genomes with little impact on microbiome. Serving as an adaptive immune system in prokaryotes, CRISPR-Cas system can recognize and degrade specific DNA targets through the guide of CRISPR RNA (crRNA)^11^. CRISPR-Cas systems are grouped into two classes and six types, among them, the type II CRISPR-Cas system, a widely used genome editing tool in eukaryotes, has been predominantly developed as antimicrobial^12–18^. However, the conventional Cas9 technology is difficult to manipulate in most prokaryotes, due to its large size and severe toxicity to host cells^19, 20^. The type I CRISPR-Cas system is more prevalent in prokaryote^21^. Type I-E CRISPR Cas system from *Escherichia coli* includes CRISPR array and the *cas* genes expressing Cas3 and five different proteins, Cse1, Cse2, Cas7, Cas5, and Cas6e (in 1, 1, 6, 1, and 2 copies, respectively) namely Cascade^22^. CRISPR array is transcribed as a long precursor CRISPR RNA (pre-crRNA) that is processed by Cas6e through cleavage at a specific site in the repeat sequence to produce the 61-nucleotide (nt) mature crRNA. Cascade, which scans and associates with the protospacer adjacent motif (PAM)^23^, facilitates recognition of DNA target by crRNA and formation of ‘R-loop’ structure^24^. Subsequently, the Cas3 effector is recruited to Cascade to degrade the target^25^. Histone H-NS (Heat-stable nucleoid-structuring protein) is a global regulator of many gram-negative bacteria and can inhibit transcription. Pul *et al* reported that H-NS can bind to the P*_cas_* and P*_crispr_* promoters of the *E. coli* type I-E CRISPR-Cas system, preventing RNA polymerase from interacting with the promoter, thereby inhibiting the transcription of the *cas* genes and pre-crRNA ^26^. As an antagonist of H-NS, LeuO can relieve the inhibitory effect of H-NS on the P*_cas_* promoter. Thus, when LeuO binds to the P*_cas_* promoter, it forms a barrier at the transcription initiation site of the *cas* gene, preventing H-NS from binding at that site, thereby activating the expression of P*_cas_*. The type I-E system has been reconstituted in *Saccharomyces cerevisiae* to prevent MGE^27^, but has rarely been used for reducing transfer of ARGs in probiotic.

For more than a century, *Escherichia coli* strain Nissle 1917 (EcN) has been used as a probiotic for treating human intestinal diseases, such as pathogen infection, diarrhea, intestinal epithelial barrier dysfunction and inflammatory bowel disease ^28–34^. Genetically engineered EcN has shown promising application as living therapeutic in treating colitis, colorectal cancer and metabolic disease ^35–41^. EcN has also been used as a vehicle for delivering the type II CRISPR-Cas systems to kill pathogens in animal models ^42^. However, strategies for reducing accumulation of ARGs in probiotics remain scarce. In this study, we used plasmids carrying antibiotic-resistant genes (ARGs), which expressed a green fluorescence gene, to precisely evaluate transfer of ARGs to EcN. Then, we directed the endogenous type I-E CRISPR-Cas system to multiple ARGs *mcr-1*, *bla*_NDM-1_ and *tet*(X) in a *E. coli* K12 strain (i.e. BW25113), and integrated the type I-E CRISPR-Cas system from BW25113 into EcN. The engineered EcN showed immunity against transfer of ARGs *mcr-1*, *bla*_NDM-1_ and *tet*(X) both *in vitro* and *in vivo*. Our work not only showed efficient transfer of ARGs to EcN in living animals, but also generated a probiotic chassis with immunity against transfer of ARGs, paving the way of clinical application of engineered EcN.

## Results

### Efficient transfer of ARGs to EcN

EcN has been developed as a vehicle for delivery of the CRISPR-Cas system^43^. But its potential risk as a reservoir of ARGs has been insufficiently evaluated. Previous studies showed that EcN received ARGs on plasmids in gut microbiome^44^. Whereas, rates of plasmid transfer into EcN vary largely, in a range from an undetectable level to 10^-1^, depending on the type of conjugative plasmid carrying ARGs, donor strain and experimental systems^45–48^. We evaluated transfer of ARGs to EcN with the donor strain *E. coli* WM3064 carrying the RP4 conjugative system, which has an extremely wide host range and can deliver DNA to a variety of microbial organisms, especially gram-negative bacteria^49, 50^. To precisely evaluate conjugative transfer rate of plasmids carrying ARGs, a reporter plasmid pGLO-J23100-*gfp*, which expressed green fluorescence protein (GFP), was transformed into EcN, yielding the recipient strain EcN-GFP (Figure 1a). WM3064 is an auxotrophic strain whose growth relies on the supplementation of diaminopimelic acid (DAP) in the medium ^51–53^. Thus, transconjugants can be easily selected by antibiotic resistance on LB plates without DAP^51^ (Figure 1a, 1b). Transfer of the conjugative plasmids pHG101 and its derivative pHG101-*tet*(X) (constructed by inserting a *tet*(X) gene into pHG101) from WM3064 to EcN-GFP was detected (Figure 1c). Frequencies of transfer of ARGs on plasmids pHG101 and pHG101-*tet*(X) were 2.44 × 10^-1^ and 1.53 × 10^-1^ respectively (Figure 1c), revealing efficient transfer of ARGs to EcN *in vitro*.

**Figure 1.**
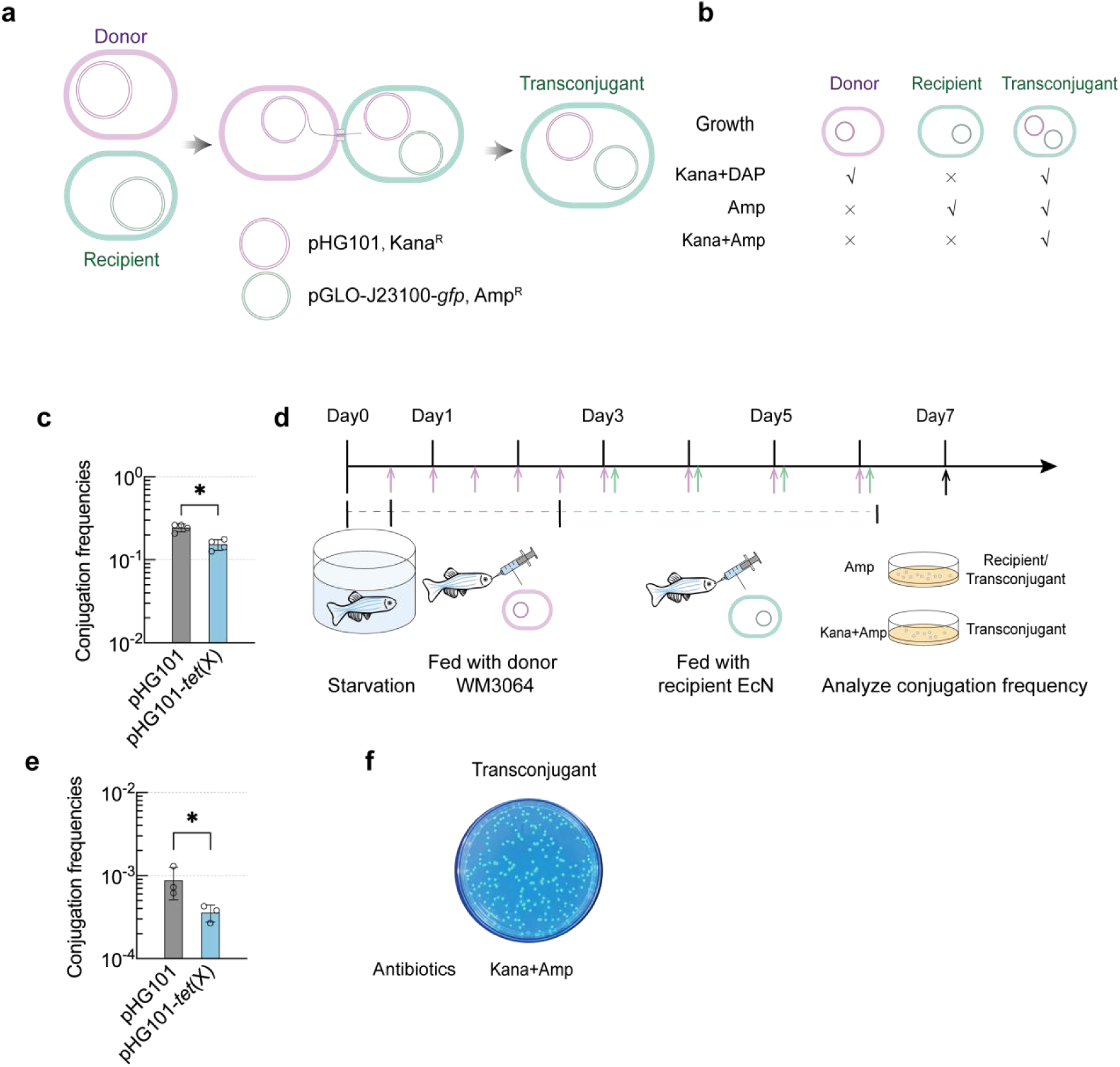
Efficient transfer of antibiotic resistance plasmids to EcN. (**a-b**) (**a**) Schematic illustration depicting transfer of conjugative plasmid pHG101 from donor WM3064 to recipient EcN. The purple oval represents the donor strain WM3064 carrying pHG101 (Kana^R^) while the green oval represents the recipient strain EcN carrying pGLO-J23100-*gfp* (Amp^R^) (EcN-GFP), or the transconjugant carrying pHG101 and pGLO-J23100-*gfp.*(**b**) The growth of auxotrophic strain WM3064 (donor) relies on the supplementation of diaminopimelic acid (DAP) in the medium containing. The antibiotic ampicillin (Amp) can be used to select transconjugants and recipients, and the combination of antibiotics ampicillin (Amp) and kanamycin (Kana) is used to select only transconjugants. (**c**) Conjugation frequency of pHG101 and pHG101-*tet*(X) transferring from WM3064 to EcN *in vitro*, pHG101-*tet*(X) (pHG101 plasmid containing the tetracycline-resistant *tet*(X) gene). (**d**) Overview of*invivo*bacterial conjugation assay in zebrafish model. Starvation-treated zebrafish were administered with WM3064 (1 × 10^6^ CFU, every 12 h for 6 times), then WM3064 and EcN were given everyday, separatively (at 2-hour intervals), subsequently the intestine was harvested and the conjugation frequency was analyzed. (**e**) Conjugation frequency of pHG101 and pHG101-*tet*(X) transferring from WM3064 to EcN in zebrafish intestine. (**f**) Colonies with green fluorescence formed by transconjugants on plates. Bright green fluorescence was clearly visible under a blue light transilluminator.

Most studies have conducted experiments showing transfer of ARGs *in vitro*^54^. As a useful model to study human intestinal pathogens^55^, zebrafish (*Danio rerio*) have been extensively used to investigate microbiome-host interactions and evaluate the toxicity of drugs or environmental pollutants^56–58^. A recent study has utilized the zebrafish intestine for testing the colonization and of therapeutic effect EcN against pathogen infection^55^. To assess the *in vivo* transfer rate of ARGs to EcN, we selected zebrafish as the model. Zebrafish were starved for 12 h, then fed with the donor strain WM3064 every 12 h for 6 times to facilitate its colonization in the intestine of the host. After feeding with WM3064, the recipient strain EcN-GFP was supplemented for 4 times every 24 h (Figure 1d). The number of recipients and transconjugants from intestines of zebrafish was detected. The conjugation frequencies of pHG101 and pHG101-*tet*(X) were 8.81 × 10^-4^ and 3.59 × 10^-4^ respectively (Figure 1e), and transconjugants with antibiotic resistance and green fluorescence were easy to detect(Figure 1f), revealing efficient transfer of ARGs on plasmids in zebrafish gut.

### Engineering CRISPR for directing the CRISPR-Cas system to cleave multiple ARGs

To direct the type I-E CRISPR-Cas system to T-MNT, a total of 9 spacers targeted to *mcr-1*, *bla*_NDM-1_ and *tet*(X) genes (three spacer sequences per target gene) were chosen and incorporated into the CRISPR array to generate CR_MNT_. The spacers were preferentially chosen to map with conserved regions of the genes to minimize the risk of escape mutants (Figure S1). To obtain a functional ARG-targeting CRISPR-Cas system, CR_MNT_ was expressed from a constitutive/strong promoter J23110. To test the feasibility of eliminating ARGs, three antibiotic resistance genes, *mcr-1*, *bla*_NDM-1_ and *tet*(X) were chosen as representative epidemic ARGs. By separately cloning these ARGs into T vector, T-*mcr-1*, T-*bla*_NDM-1_ and T-*tet*(X) (GenBank accession numbers PP555434, PP555435 and PP555436) (T-MNT) were generated as ARG-carrying plasmids, which can be targeted by the engineered CRISPR-Cas system. T-MNT plasmids were transformed into *E. coli* K12 strain BW25113, which harbors a type I-E CRISPR-Cas system, and the clearance of T-MNT was evaluated by calculating their transformation efficiencies in the BW25113 expressing CR_MNT_.

LeuO and H-NS are known strong regulators of CRISPR immunity ^26, 59^, H-NS bind to and silence P*_cas_* to suppress the transcription of *cas* genes, while LeuO activates *cas* genes transcription by relieving transcriptional suppression by H-NS. For activating CRISPR-Cas, *leuO* gene expressed from an arabinose-inducible promoter P_BAD_ was introduced into the CR_MNT_ plasmid to generate LeuO-CR_MNT_ (Figure 2a). Compared with cells carrying non-activated CRISPR (Vector), the transformation efficiencies of T-MNT were reduced by 96.8%, 97.5% and 94.4% respectively in cells with activated CRISPR (LeuO-CR_MNT_, 30mM) (Figure 2b), showing a remarkable decrease of transformation efficiencies of all the three CRISPR-targeted ARG carrying plasmids. The leaky expression of LeuO was sufficient to achieve effective cleavage by CRISPR-Cas, as the transformation efficiencies of T-MNT were reduced by 93.1%, 95.9% and 82.5% without arabinose (Figure 2c). *hns* inactivation (Δ*hns*) result in stronger CRISPR immunity against ARGs than over-expressing LeuO, as the transformation efficiencies of T-MNT were reduced by 99.6%, 99.8% and 99.8% (Figure 2d), suggesting the robust clearance activity of ARGs by type I-E CRISPR-Cas system. Together, our results show that the type I-E CRISPR-Cas system can be directed to cleave multiple ARGs in BW25113.

**Figure 2.**
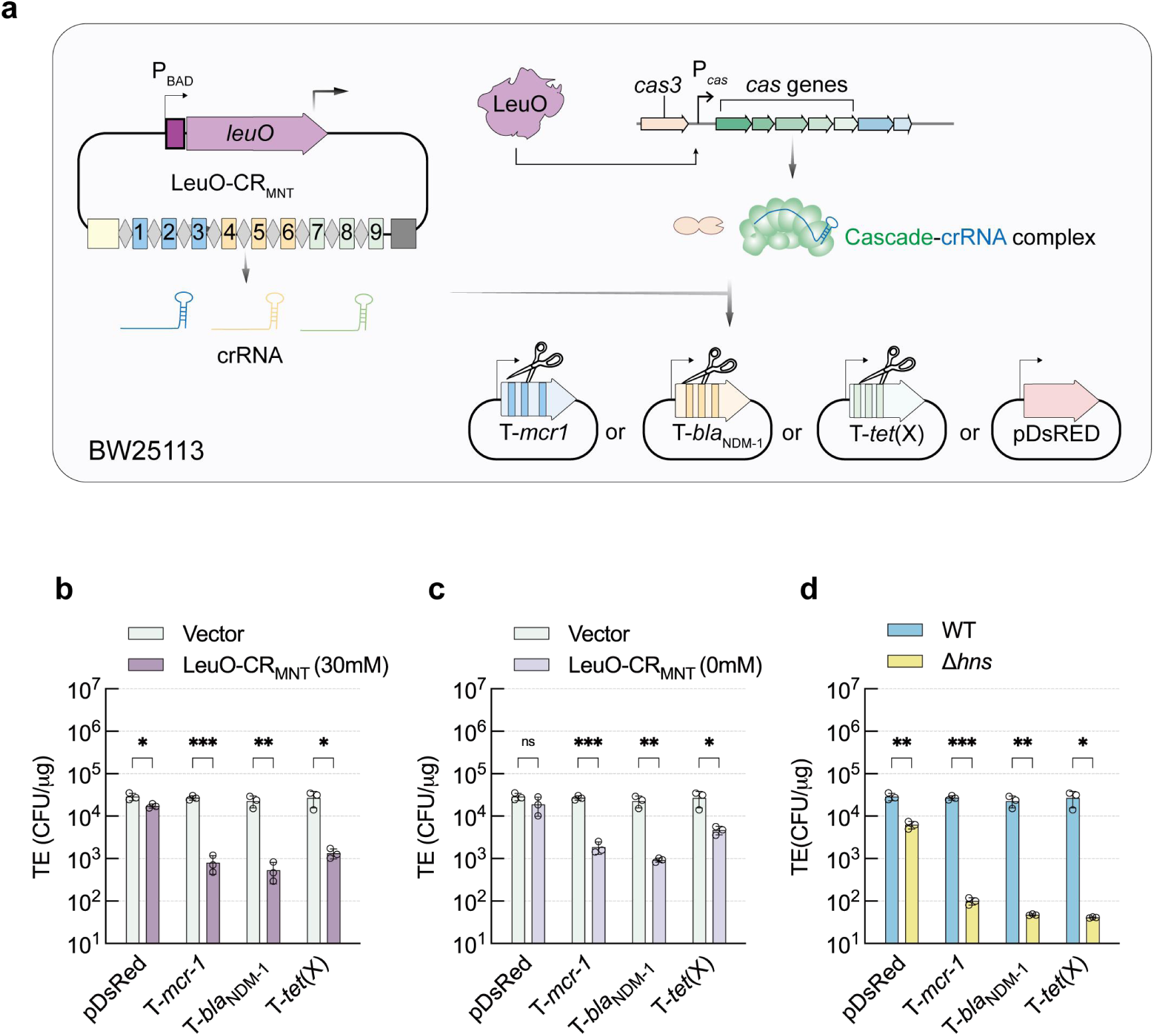
Directing the endogenous type I-E CRISPR-Cas system to cleave ARGs in *E.coli*BW25113. (**a**) Schematic diagram of directing the type I-E CRISPR-Cas system in BW25113 to target the antimicrobial resistance (AMR) plasmids (T-*mcr1*, T-*bla*_NDM-1_ and T-*tet*(X)). A CRISPR-nontargeted plasmid pDsRED was used as a negative control. The plasmid (LeuO-CR_MNT_) was used to overexpress the P*_cas_* activator LeuO from the P_BAD_ promoter and the targeting crRNA (CR_MNT_) from J23100 promoter. CR_MNT_ was created by inserting 9 spacers (3 each mapping *mcr1*(blue rectangles 1-3), *bla*_NDM-1_ (yellow rectangles 4-6) and *tet*(X) (green rectangles 7-9) into a CRISPR array under the control of J23110 promoter (yellow box), with a terminator (grey box) preventing read-through. (**b-c**) Transformation efficiency (TE) of pDsRED or the AMR plasmids (T-*mcr1*, T-*bla*_NDM-1_ and T-*tet*(X)) in CRISPR-Cas activated cells (expressing LeuO and CR_MNT_) or CRISPR-Cas silenced cells (expressing only CR_MNT_, Vector). Cells were treated with (**b**) or without (**c**) 30mM arabinose for induction of LeuO. (**d**) Transformation efficiency (TE) of pDsRED or the AMR plasmids (T-*mcr1*, T-*bla*_NDM-1_ and T-*tet*(X)) in WT or *hns*mutant cells.

### Reconstruction of an inducible type I-E CRISPR-Cas system on plasmid

Since no *cas* genes and CRISPR arrays have been found on the genome of EcN, the type I-E CRISPR-Cas system from *E. coli* K12 were selected for designing a programmable anti-ARG system in EcN. To this end, components of the type I-E CRISPR-Cas system were cloned from *E. coli* BW25113 and inserted into different plasmids (Figure 3a). The pSU-*cas* plasmid, expressing Cascade proteins, consists of five cassettes with different *cas* genes (*cse1*, *cse2*, *cas7*, *cas5*, *cas6e*) under the control of the arabinose-inducible promoter P_BAD_. For the expression of the Cas3 nuclease, we designed the pSU-*cas3* plasmid by cloning *cas3* under the P_BAD_ promoter (Figure 3a, 3b). To validate correct construction of plasmids pSU-*cas3* and pSU-*cas*, the two plasmids were transformed into a *cas3*-deleted strain BW25113 Δ*hns* Δ*cas3* and a *cas*-operon-deleted strain BW25113 Δ*hns* Δ*cas* respectively (Figure 3a, 3b) and the CRISPR-Cas immunity were evaluated in complemented mutants. Transformation efficiency of the CRISPR-targeted plasmid pCRI was examined in BW25113 Δ*hns* Δ*cas3* strain carrying pSU-*cas3* and BW25113 Δ*hns* Δ*cas* strain carrying pSU-*cas*. The transformation efficiency of pCRI was more than 100-fold lower in Δ*hns* Δ*cas3* expressing pSU-*cas3* comparing with that carrying the empty vector pSU19. The similar effect was observed in Δ*hns* Δ*cas* expressing pSU-*cas* (Figure 3c, 3d). In contrast, transformation efficiency of the non-targeted plasmid pDsRED was not affected in Δ*hns* Δ*cas3* and Δ*hns* Δ*cas* carrying the empty vector. Interestingly, strong immunity against pCRI of the two strains against the targeted plasmid were also observed without induction by arabinose, implicating that leaky expression of either Cas3 or Cascade was sufficient for establishing immunity against CRISPR-targeted plasmid (Figure 3c, 3d).

**Figure 3.**
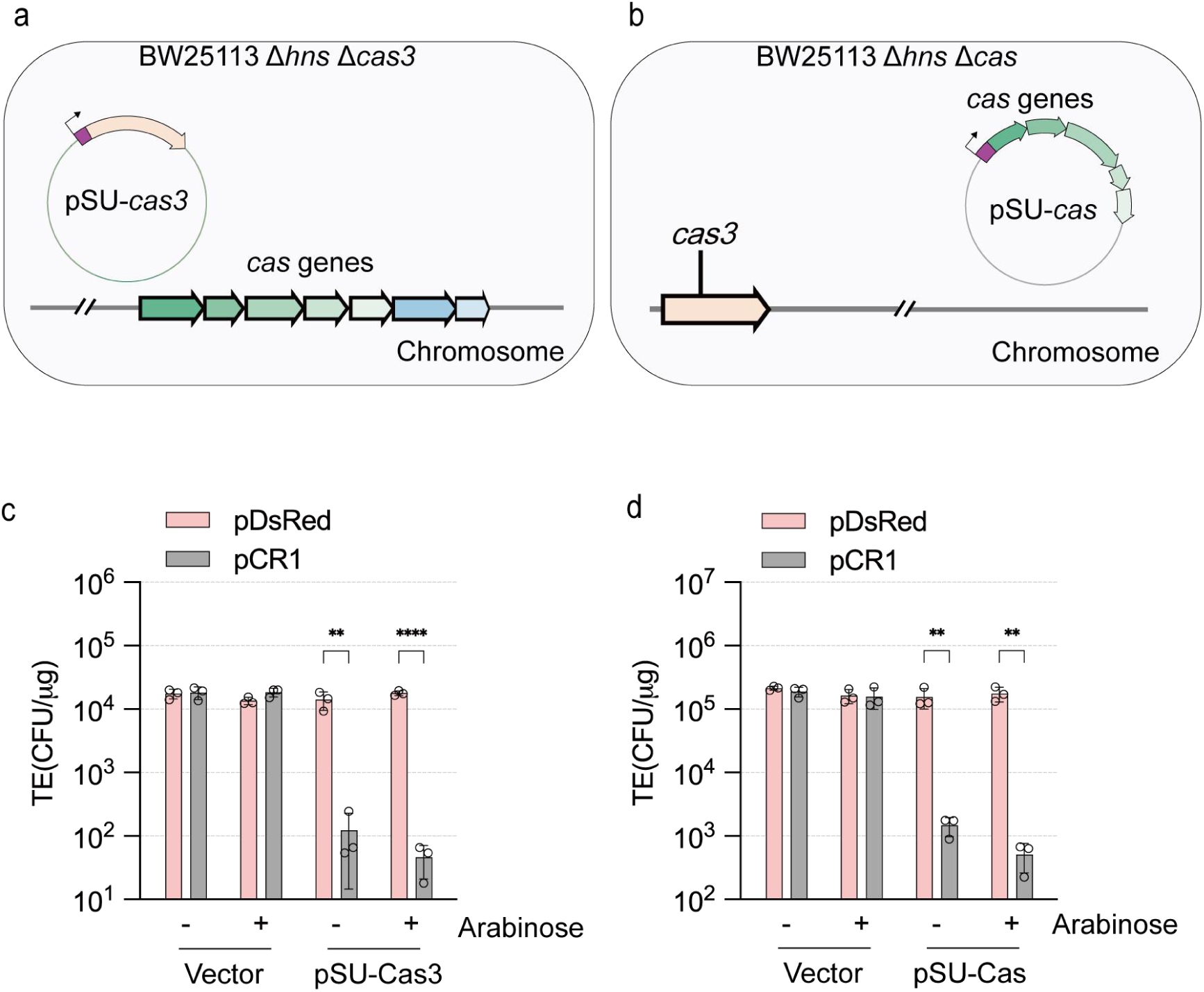
Reconstitution of an inducible type I-E CRISPR-Cas system on plasmid in EcN. **(a-b)** Design of the arabinose-induced type I-E CRISPR-Cas system on plasmid. By expressing *cas3*or the *cas*genes from arabinose-inducible promoter (P_BAD_) on plasmids pSU-*cas3*(**a**) or pSU-*cas*(**b**) respectively, the reconstituted CRISPR-Cas system can be guided to degrade of the corresponding target DNA. pCRI contains the CRISPR-targeted sequence, while the non-targeted plasmid pDsRED serves as a control. **(c)** Transformation efficiency (TE) of pDsRed and pCRI in BW25113 Δ*hns*Δ*cas3*containing control plasmid pSU19 (Vector) and pSU-*cas3*. Cells were treated with (+) or without (-) 30 mM arabinose for induction of *cas3*. **(d)** Transformation efficiency (TE) of pDsRed and pCRI in BW25113 Δ*hns*Δ*cas*containing control plasmid pSU19 (Vector) and *cas*genes expressing plasmids (pSU-*cas*). Cells were treated with (+) or without (-) 30 mM arabinose for induction of *cas*genes.

### Engineering EcN for cleaving ARGs with the type I-E CRISPR-Cas system

To generate a type I-E CRISPR-Cas-armed EcN strain for cleaving ARGs, CR_MNT_ that expressed crRNAs targeting to *mcr-1*, *bla*_NDM-1_ and *tet*(X) was inserted into pGLO-*cas3* (Figure 4a). The generated recombinant plasmid (pGLO-*cas3*-CR_MNT_) and pSU-*cas* were transformed into EcN, yielding the engineered strain EcN-V1 (Figure 4a). To mimic the HGT of ARGs through conjugation, the conjugative plasmids pHG101 carrying *mcr*-1, *bla*_NDM-1_ or *tet*(X) were transferred from WM3064 (donor strain) to EcN-V1 (recipient strain). EcN carrying pSU19-*cas* and pGLO-*cas3* (EcN-C1, without CR_MNT_) was used as a negative control of recipient strain. (Figure 4a). The conjugation frequency of pHG101-*mcr-1* was 349 folds lower in EcN-V1, with regard to that in EcN-C1 (Figure 4b). Whereas, we only observed about 3.75-fold decrease of conjugation frequency of pHG101-*bla*_NDM-1_ (Figure 4b), and 23-fold decrease of the conjugation frequency with pHG101-*tet*(X) (Figure 4b). In contrast, the conjugation frequencies of non-targeted plasmid pHG101 were similar in EcN and EcN-V1. (Figure 4b). We found the cleavage of pHG101-*mcr-1* was stronger in EcN than that in the CRISPR-Cas activated BW25113 (Figure 4b), while the cleavage of pHG101-*bla*_NDM-1_ was attenuated in EcN, indicating the CRISPR-Cas-armed EcN was effective to limit the uptake of plasmids, and the cleavage preference was different with BW25113.

**Figure 4.**
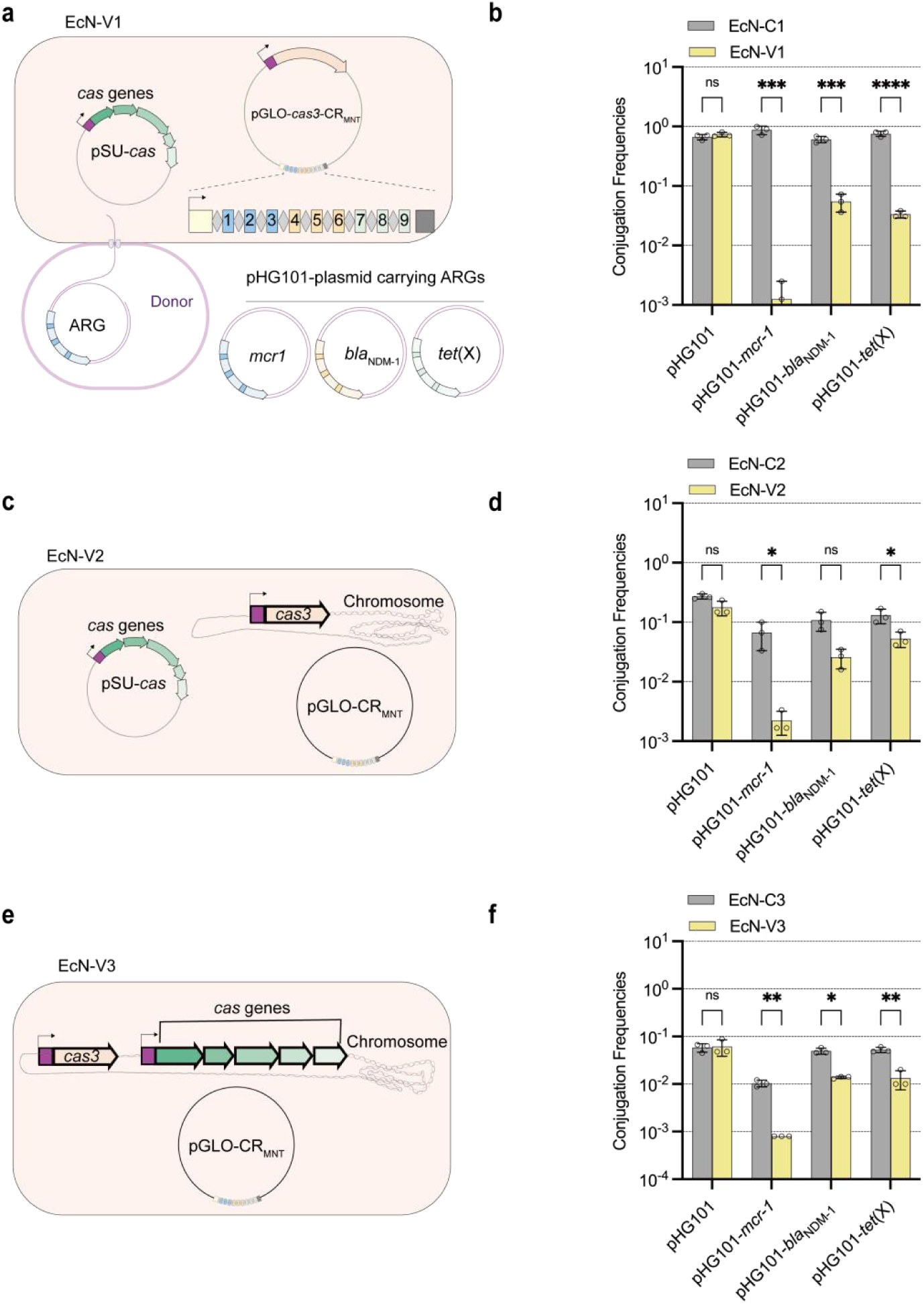
Eliminating ARGs by expressing the type I-E CRISPR-Cas system in EcN. (**a**) Schematic representation of the EcN-V1 cells, which carry pSU-*cas* expressing *cas* genes and pGLO-*cas3*-CR_MNT_ plasmid expressing *cas3* and crRNA targeting to *mcr-1*, *bla*_NDM-1_ and *tet*(X). For the negative control EcN-C1 cells, the pGLO-*cas3* plasmid containing no CRISPR array was used instead of pGLO-*cas3*-CR_MNT_. Plasmid ARG (pHG101-*mcr-1*, pHG101-*bla*_NDM-1_, pHG101-*tet*(X)) or empty vector pHG101 were transferred from WM3064 to EcN V1 or EcN-C1. **(b)** Plasmid elimination measured by conjugation frequencies of pHG101-*mcr-1*, pHG101-*bla*_NDM-1_, pHG101-*tet*(X) or empty vector pHG101 from WM3064 towards recipient cells EcN V1 or EcN-C1. (**c**) Schematic representation of the EcN-V2 cells. *cas3*expressed from an arabinose-induced promoter (P_BAD_, shown as a dark purple box) was integrated in the genome, while *cas*genes was expressed from pSU-*cas*, and crRNA expressed from the plasmid pGLO-CR_MNT_. For the negative control EcN-C2 cells, the empty vector pGLO-empty was used instead of pGLO-CR_MNT._ **(d)** Plasmid elimination measured by conjugation frequencies of pHG101-*mcr-1*, pHG101-*bla*_NDM-1_, pHG101-*tet*(X) or empty vector pHG101 from WM3064 towards recipient cells EcN V2 or EcN-C2. (**e**) In the EcN-V3 cells. The *cas*genes and *cas3* were all integrated in genome, while crRNA was expressed from the plasmid pGLO-CR_MNT_. For the negative control EcN-C3 cells, the empty vector pGLO-empty was used instead of pGLO-CR_MNT_. (**f)** Plasmid elimination measured by conjugation frequencies of pHG101-*mcr-1*, pHG101-*bla*_NDM-1_, pHG101-*tet*(X) or empty vector pHG101 from WM3064 towards recipient cells EcN V3 or EcN-C3.

Previous study showed ectopic over-expression of Cas3 in *E. coli* cells could stimulate uncontrolled replication of plasmids with ColE1 replicons^60^. In our study, spacers targeted to *tet*(X) were found mutated frequently on pGLO-*cas3*-CR_MNT_ (data not shown), suggesting a side effect caused by overexpressing Cas3. To avoid unexpected mutation and plasmid instability, we integrated *cas3* into the genome of EcN, which was then transformed with pSU-*cas* and pGLO-CR_MNT_ for generating a complete ARG-targeting type I-E CRISPR-Cas system in EcN (named EcN-V2) (Figure 4c), EcN-C2 was the same with EcN-V2 except carrying pGLO empty vector instead of pGLO-*cas3*-CR_MNT_. Comparing with EcN-C2, EcN-V2 was able to restrict the transfer of ARGs. The conjugation frequency of pHG101-*mcr*-1, pHG101-*bla*_NDM-1_ and pHG101-*tet*(X) were reduced by 30, 4.2 and 2.49-fold respectively (Figure 4d). Although the amount of *cas3* expressed from genome was less than that from an inducible promoter from plasmid, EcN-V2 also effectively limit the transfer of the ARGs. To avoid loss of the large-size plasmid pSU-*cas*, *cas* genes were further integrated into EcN-V2 genome to generate EcN-V3 (Figure 4e). Also, the negative control strain EcN-C3 was the same with EcN-V3 except carrying pGLO empty vector instead of pGLO-CR_MNT_. When using EcN-V3 as a recipient strain, the conjugation frequency of pHG101-*mcr*-1, pHG101- *bla*_NDM-1_ and pHG101-*tet*(X) decreased by 57.8, 3.6, and 4-fold, respectively, compared with the EcN-C3. (Figure 4f). These results demonstrated that our CRISPR-Cas-armed EcN can serve as a chassis with high resistance to the invasion of ARGs.

### EcN expressing Cas3 shows growth advantage over that expressing Cas9

Despite its extensive application for genome editing in eukaryotes^61^, previous study showed type II nuclease Cas9 can efficiently block transcription initiation and inhibit the growth of *E. coli* ^62^, suggesting severe toxicity to bacteria. Thus CRISPR-Cas9 is supposed to be challenging to exploit in prokaryotes. Therefore the type I-E nuclease Cas3 from *E.coli* can be highly suitable for expressing in EcN. To test this, we co-incubated EcN expressing the arabinose-inducible Cas3 (EcN-I) or Cas9 (EcN-II) in the same LB culture (Figure 5a). EcN-I exhibits increasing growth advantage over EcN-II in 36 h. The advantage is more significant when inducing Cas3 or Cas9 by addition of 30 mM arabinose, the proportion of EcN-I reaches 99% at 12 h and maintains at a level over 95% (Figure 5a), indicating EcN expressing Cas3 has better adaptive capability than EcN expressing Cas9.

**Figure 5.**
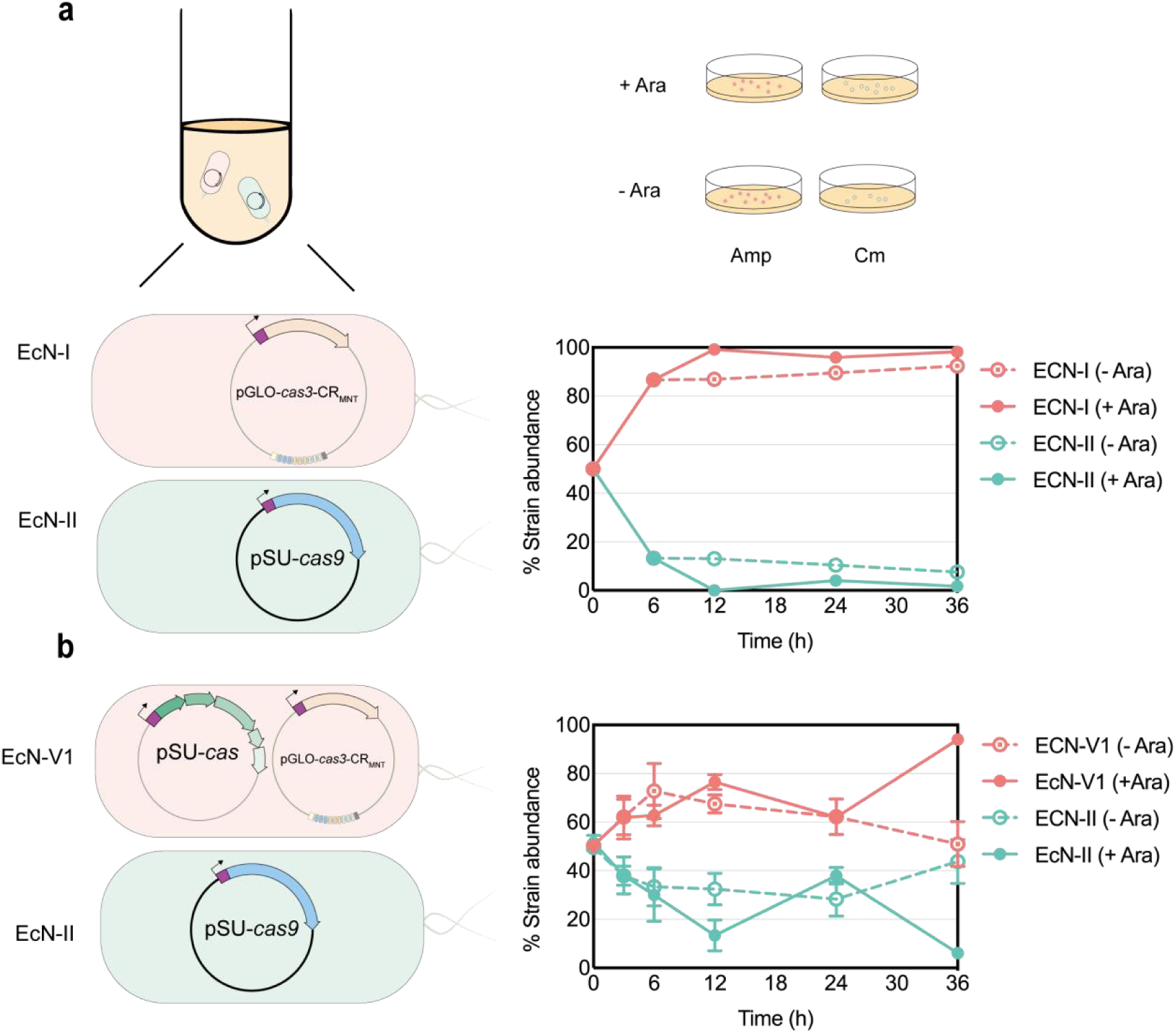
Competitive experiment of EcNs expressed type I and type II CRISPR-Cas System. (**a**) Growth competition of EcN-I and EcN-II. Equal amount of *E.coli*derivatives (E-I, E-II) strains were mixed and inoculated into antibiotics-free LB medium with or without 30 mM arabinose, respectively. The cell culture was incubated with shaking at 37℃ for 36 h. The percentage of EcN-I (red), EcN-II (green) live cells were measured by counting colonies on plates containing different antibiotics. The growth of EcN-I (red), EcN-II (green) supplied with arabinose (+ Ara) are indicated by solid lines, while EcN-I (red), EcN-II (green) supplied without induction(-Ara) is indicated by dashed lines. **(b)** Growth competition analysis of EcN derivatives in simulated mouse intestinal environments. Equal amount of *E.coli*derivatives (EcN-V1, EcN-II) were mixed and inoculated into 50 mL LB medium containing 0.2 g mouse feces with (solid lines, + Ara) or without (dashed lines,-Ara) 30 mM arabinose and grown at 37℃. The percentage of EcN-V1 (red), EcN-II (green) live cells were measured by counting colonies on plates containing different antibiotics.

The type I-E CRISPR-Cas immunity requires Cas3 and multisubunit protein-RNA complex Cascade. To test the adaptive capability in EcN with all the components for type I-E CRISPR-Cas immunity, the EcN-V1 strain carrying the plasmid pSU-*cas* and pGLO-*cas3*-CR_MNT_, and EcN-II carrying the plasmid pSU-*cas9* was used. Competitiveness of EcN-V1 and EcN-II was compared in the culture mimicking mouse gut microbiota by adding mouse feces to LB^63^. When no arabinose (inducer) was added, the proportion of EcN-V1 was higher than E-II at 6 h but the abundance of these two strains became gradually consistent during the next 30 h. While in the presence of inducer, EcN-V1 showed a significant growth advantage than type II CRISPR-Cas system at 36 h (Figure 5b). These results indicate that the type I-E CRISPR-Cas3 system can be more adapted for EcN than the type II CRISPR-Cas9 system, the former can lead to better application prospects *in vivo*.

### Engineered EcN can restrict transfer of ARGs *in vivo*

Although genomic expression of Cas3 and Cascade (EcN-V3) significantly decreased the transfer rates of ARGs in LB culture medium (Figure 4f), whether EcN-V3 can colonize, adapt and show immunity against ARGs in the gut is not tested. To this end, we performed *in vivo* conjugation of EcN-V3 to further assess its ability of blocking transfer of ARGs in the zebrafish gut microbiota. The donor WM3064 and recipient EcN-C3 or EcN-V3 were administrated into zebrafish by gavage. After 7 days treatment, the bacteria were extracted from zebrafish gut, and the conjugation frequencies of pHG101 and its derivative plasmids were measured (Figure 6a and 6b). As expected, the conjugation frequency of pHG101-*mcr*-1, pHG101-*bla*_NDM-1_, pHG101-*tet*(X) was reduced by 15.3, 5.16 and 29.88-fold respectively in EcN-V3 compared with that in EcN-C3 (Figure 6a), confirming efficient cleavage of ARGs in the engineered EcN in zebrafish gut.

**Figure 6.**
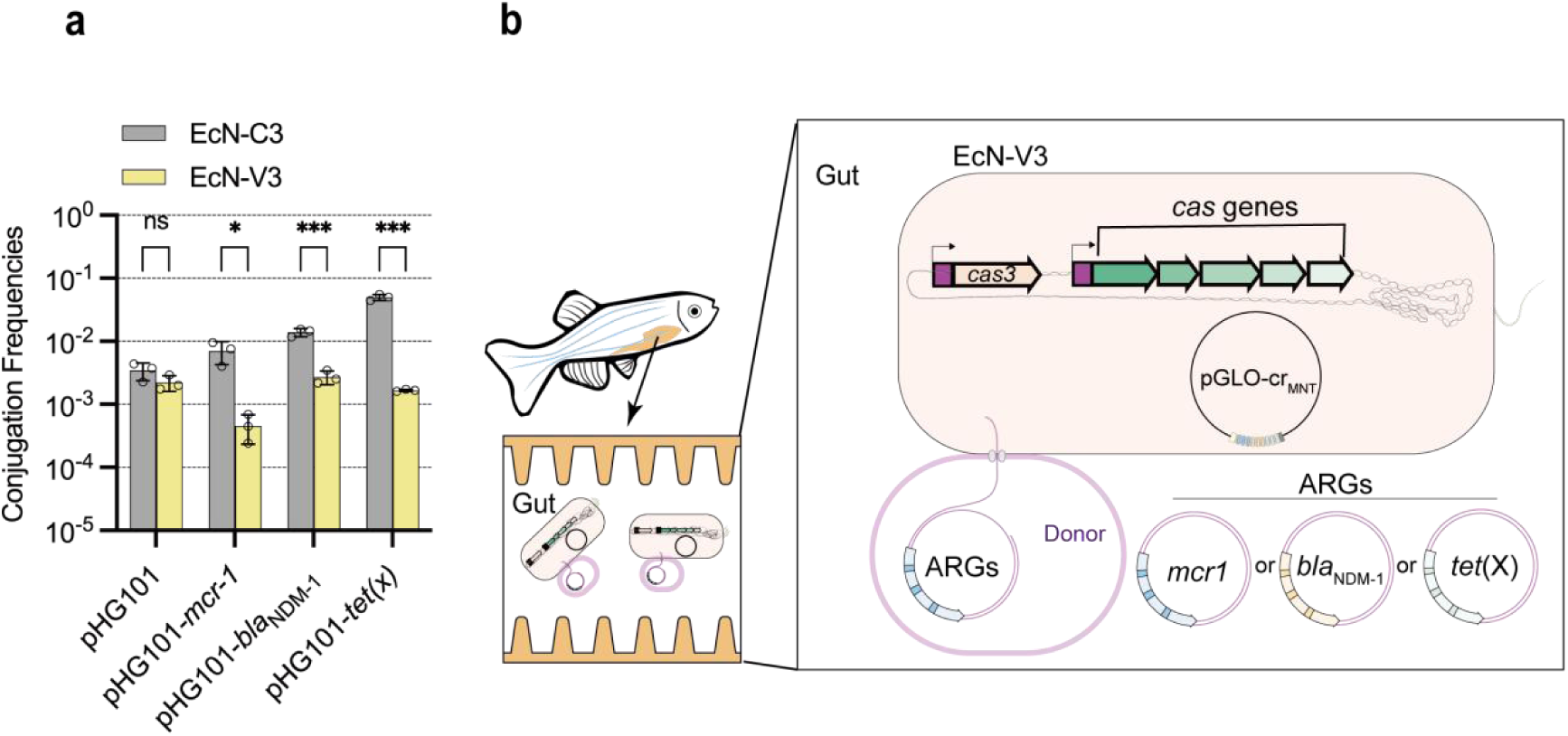
Effects of engineered EcN on restricting ARGs *invivo*. (**a**) Engineered EcN limits the transfer rates of ARGs in zebrafish intestine. Zebrafish were treated with WM3064 carrying pHG101-*mcr-1*, pHG101-*bla*_NDM-1_, pHG101-*tet*(X) or empty vector pHG101, and then EcN-V3 or EcN-C3. *Invivo*plasmid ARGs elimination of ARGs plasmids in zebrafish gut were measured by calculating conjugation frequencies. (**b**) Conjugation frequencies of pHG101-*mcr-1*, pHG-101-*bla_NDM_*, pHG101-*tet*(x) or empty vector pHG101 transferred from WM3064 to recipient cells EcN-V3 or EcN-C3 in zebrafish intestine

## Discussion

Despite rapid development and spread of MDR due to antibiotic abuse, strategies for slowing down this trend remain scarce. Probiotics, bacteriophages and CRISPR-Cas antimicrobials have been most intensively investigated as alternatives of antibiotics ^63–65^. Delivering CRISPR-Cas systems by probiotics such as engineered EcN through conjugation is promising, because this strategy does not trigger host response that could otherwise activate the endogenous CRISPR-Cas system for combating phage infection^66^. Besides, although probiotics have also been developed as living therapeutics for treating pathogen infection, metabolic diseases, inflammatory bowel disease and cancer^31–35, 37–41^, it remains obscure whether living therapeutics predispose to accumulate and spread ARGs in gut microbiome, exacerbating MDR. In this study, we showed that conjugative plasmids can be transferred into a probiotic strain EcN with high efficiency both *in vitro* and *in vivo*, providing additional experimental evidence showing the risk of a probiotic strain in accumulating ARGs. More importantly, we integrated a type I-E CRISPR-Cas system from *E. coli* BW25113 into EcN, endowing this strain with immunity against multiple ARGs. We further showed that the engineered EcN not only successfully blocked the entry of multiple ARGs *in vitro*, but also limited transfer of them in an animal model (i.e. zebrafish). Particularly, because targeted ARGs can be recognized by different crRNAs in our engineered EcN, it would be convenient to expand its immunity to other ARGs by simply adding new spacers in the CRISPR array. Our work would facilitate the clinical application of EcN as a chassis for the development of the next generation of living therapeutics with reduced risk of accumulation of ARGs.

Few studies have analyzed the efficiency of horizontal transfer of AMR *in vivo*. Transformation or conjugation efficiencies were conventionally measured by calculating colonies on plates with appropriate antibiotics^63, 67^. In the past few years, fluorescent or bioluminescent reporters are extremely useful to tag live biotherapeutic bacterial strains during *invivo* preclinical assays^68, 69^. By taking advantage of GFP reporters in EcN, we are able to visualize and distinguish bacteria strains accurately from animal model (Figure 1f). By combining the GFP reporter with flow cytometry, future applications in analyzing samples in bulk would be convenient.

Previous work showed that a conjugative endogenous CRISPR-Cas system was engineered for curing high-risk IncFII plasmids in *Klebsiella pneumoniae*^67^. In our work, type I-E CRISPR-Cas system was engineered into the probiotics EcN to limit spreading of ARGs. As a traditional probiotic with a well-established safety record, EcN is wildly used in treating enteric infections and various inflammatory disorders^33, 34, 38^. Our CRISPR-Cas3-armed EcN exhibits advantages in clinical application prospects over the other pathogenic bacteria since the later contains virulence genes. Moreover, EcN is now widely utilized in the bioindustry for producing metabolites of industrial interest ^70–75^. Harmful phages or genes (e.g., ARGs) were potentially accumulated and spread in laboratory and industrial-engineered EcN. Our CRISPR-Cas-armed EcN endows this microbial chassis restriction for high-risk HGT to further improve the safety of EcN-based therapeutics or biosynthesis. The work in *K.pneumoniae*reported a high plasmid curing efficiency *in vitro* (8-log decrease) and *in vivo* (∼100% curing) in a *Galleria mellonella* infection model. In our work, CRISPR-Cas3 armed EcN blocks the transfer of multiple ARGs, with a plasmid curing efficiency of 349 folds (*mcr-1*) *in vitro* and 5.16∼29.4-fold *in vivo* (zebrafish model). Even higher cleavage efficiencies may be achieved by further improving EcoCas3’s activity through structure-guided engineering or directed evolution, or regulating the activator (e.g. LeuO) to activate the expression of type I CRISPR gene in cells. In our work, *cas3* and *cas* genes were integrated into genome of EcN (EcN-V3) to increase genetic stability of CRISPR-Cas system and reduce the addition of antibiotics to maintain plasmid stability. EcN-V3 can also be adapted to eliminate specific ARGs or phages when combined with different crRNAs.

Our work takes advantage of the type I-E CRISPR-Cas system to achieve complete cleavage of multiple ARGs. The CRISPR-Cas9 tool has been broadly engineered to restrict antibiotic-resistant bacteria^15^. However, deletions of ARGs from these two types of CRISPR-Cas are of different nature. Cas9 generates single blunt cut^76^, besides it may cause a few indels (hundred nucleotides), rare distal deletions and complex chromosomal rearrangements around the target site in mESCs^77^. Cas3 causes unidirectional cleavage sites accompanied by small indel formation, which extensively destroy the entire target DNA from upstream of the PAM^25, 78^. A high efficiency of ARGs degradation can be achieved from the cleavage pattern of Cas3 because the cleaved short fragments can scarcely be repaired. Similarly, the type I CRISPR-Cas systems have been used in human cells to introduce large genomic deletions at specific locations^79–81^, and in *Saccharomyces cerevisiae* to target plasmids^27^. Our work also showed growth advantage of EcN expressing Cas3-Cascade over that expressing Cas9. Although the type II CRISPR-Cas9 system is toxic to many host bacteria, previous work applied this system in killing pathogens or eliminating ARGs^15, 17, 20, 63^. Whereas, an excellent probiotic should not only exert a beneficial effect, but also out-compete other bacteria in the complex ecosystem of the gut microbiome. Well-growth of EcN carrying the CRISPR-Cas3 system implies that our constructed ARG-restricting EcN strain should be able to better colonize and compete with the other gut microbiota in intestine. Besides, the CRISPR-Cas3 system is more precise than the CRISPR-Cas9 system because the former requires a longer target-binding spacer sequence (32-nt), and off-targeting for type I CRISPR-Cas can be further suppressed at the Cas3 recruitment step by a large conformational change in Cascade upon full R-loop formation ^82, 83^. Thus, cleavage of ARGs by our engineered type I CRISPR-Cas system can be more stringent.

Our work provides new clues for investigating the working mechanism of the type I-E CRISPR-Cas system. Although an arabinose-inducible promoter was used for controlling expression of Cascade and Cas3, we observed that their leaky expression was sufficient for activation of the type I-E CRISPR-Cas system in both BW25113 and EcN. Whereas, the effect of blocking transfer ARGs was much stronger (>10 folds) in BW25113 than that in EcN. Improving expression of Cascade and/or Cas3 by adding the inducer (i.e. arabinose) only slightly (if any) promoted the DNA cleavage activity in EcN. It is tempting to speculate the amount of Cas3, Cascade and crRNA was sophisticatedly regulated to achieve optimized activity. Previous studies have revealed the stoicheiometry of Cascade and crRNA affect CRISPR-targeted DNA cleavage. The Cas6 family members, *Thermus thermophilus* Cas6A and Cas6B (type I-E) and of *Pseudomonas aeruginosa* Csy4 (type I-F) react stoichiometrically during pre-crRNA cleavage (at an enzyme: RNA molar ratio of 2:1)^84, 85^, since these Cas6 enzymes retain tight binding for crRNA and functions as a single-turnover catalyst. While P_BAD_ promoter is moderate in EcN (unpublished data), expression of Cascade might be insufficient. The P_BAD_ promoter could be changed to a stronger one (such as P*_tac_*) to optimize the expression of Cascade and/or Cas3. Another possibility is that the activity of the type I-E CRISPR-Cas system was limited because unknown CRISPR-Cas activators were absent in EcN. Identification of these activators would provide new insights into the working mechanism of the type I-E CRISPR-Cas system in its natural bacterial host. The sophisticated regulation of type I-E CRISPR-Cas expression might be responsible for preventing excess components released into the cytoplasm, which can lead to auto-immunity. Of note, while we were constructing a plasmid expressing both Cas3 and crRNA, we repeatedly observed that spacers within the CRISPR array were disposed to be mutated, under the condition that transcription of the *cas* operon was not induced. This abnormal phenomenon might implicate that Cas3 could affect spacers within the CRISPR array without the assistance of Cascade. Parallel analysis of the type I-E CRISPR-Cas system in closely related natural and non-natural bacterial hosts would provide new perspectives on its working mechanism.

In conclusion, our work successfully constructed a type I-E CRISPR-Cas system armed EcN which can restrict transfer of multiple ARGs. We also showed that the engineered EcN was able to block transfer of ARGs in the intestine of zebrafish. This work would reduce the risk of spreading ARGs when using EcN as a living therapeutic. The strategy for reducing accumulation of ARGs in EcN could also be applied in other probiotics, preventing spread of ARGs in gut microbiome. Moreover, our engineered EcN could also be used for precise genome editing and gene expression regulation, further expanding the application scope of this strain, and also expand the application range of the CRISPR-Cas3 toolbox in the future.

## Materials & Methods

### Strains, plasmids, and growth conditions

All strains and plasmids used in this study are listed in Table 1 and Table 2 (SUPPLEMENTAL MATERIALS AND METHODS) respectively. By using a λ-RED recombination system expressed by the temperature-sensitive plasmid pKD46^86^, each component of the CRISPR-Cas system was integrated into the genome of Nissle 1917. Construction of plasmids and strains was described in the SUPPLEMENTAL MATERIALS AND METHODS. The donor strain for conjugation *E. coli* WM3064 and the mobilizable plasmid pHG101 were provided by Dr. Yin of Zhejiang University of Technology. WM3064 was grown in LB medium with lower salt concentration (10 g L^-1^ tryptone, 5 g L^-1^ yeast extract and 5 g L^-1^ NaCl) containing 0.3 mM diaminopimelic acid (DAP). Other *E. coli* strains were grown in LB medium (10g L^-1^ tryptone, 5 g L^-1^ yeast extract and 10 g L^-1^ NaCl) or on LB-agar plates. When necessary, appropriate antibiotics were supplemented at the following final concentrations: ampicillin (100 µg mL^−1^), chloramphenicol (25 µg mL^−1^) and kanamycin (50 µg mL^−1^).

### DNA manipulations

All primers used in this work are shown in Table 3 (SUPPLEMENTAL MATERIALS AND METHODS). The plasmid miniprep kit (Axygen Biotech Co. Ltd) was used for plasmid extraction. Fragments obtained from genome or plasmids were amplified using Phanta mix (Vazyme Co. Ltd), and were purified by PCR cleanup kit (Axygen Biotech Co. Ltd). The PCR product was treated by *Dpn*Ⅰ to remove the template plasmid. DNA Sequencing was performed at the Tsingke Biotechnology Co. Ltd. The CRISPR array CR_MNT_ were synthesized by Tsingke Biotechnology Co. Ltd

### Plasmid Transformation

For Figure 2b-d and Figure 3c-d, CRISPR immunity was measured by cleavage of targeted plasmid during chemical transformation. The chemical competent cells were prepared with the CaCl_2_ solution. In brief, *E. coli* strains were grown overnight in 5 mL LB at 37°C. 1 mL of the overnight culture was inoculated into 50 mL LB, and grown to an OD_600_ of 0.4 ∼ 0.6, followed *by* harvesting on ice, and washing with 100 mM CaCl_2_ solution in triplicate. The competent cells were resuspended in 100 mM CaCl_2_ solution supplemented with 10% glycerol. For Chemical transformation, 50 μL of the competent cells were mixed with 500 ng DNA and incubated on ice for 30 min. The mixture was heated at 42°C for 90 s and then placed on ice for 2 min, followed by the addition of preheated LB medium up to 1 mL. Cells were incubated at 30°C for 1 h before being spread on appropriate selective plates. The plates were cultured at 37°C for 12 ∼ 24 h before counting the number of transformants.

### In vitro bacterial conjugation assay

The donor (WM3064) and the recipient (EcN) strains were separately grown in LB with shaking at 37°C. Overnight grown cultures were inoculated into fresh LB medium at a ratio of 1:100, and incubated to an OD_600_ of 0.4-0.6 at 37°C. 2 mL of WM3064 and 1 mL EcN cells were collected by centrifugation for 3 min at 4000 rpm, followed by washing twice with antibiotic-free LB. After the last washing, the precipitate (mixture of the donor and recipient cells) was resuspended in 100 μL LB and dropped on LB agar plates supplemented with DAP, followed by incubation for 4 h at 37°C. Cells were recovered from the plate and resuspended in 1 ml LB, followed by serial dilution and plating on LB agar plates supplemented with appropriate antibiotics. The conjugation frequency was calculated by measuring the ratio of transconjugants to recipients.

### In vivo bacterial conjugation assay

Zebrafish (*Danio rerio*) of mixed sex were raised in a circulatory system at 28°C, and fed with pellet diet twice a day. To evaluate *in vivo* transfer of ARGs, Zebrafish were starved for 12 h and anesthetized on ice, before being fed with donor and/or recipient bacteria (∼1 × 10^6^ CFU) for 7 days. Then, zebrafish were dissected aseptically with sharp-topped tweezers and surgical scissors, and the entire intestine were placed in a sterile 1.5-ml centrifuge tube on ice, suspended with 50 μL PBS solution, and grinded with 3 mm zirconia magnetic beads. The suspended intestinal contents were 10-fold serial diluted, and spread onto LB agar plates supplemented with appropriate antibiotics. The conjugation frequency was calculated by measuring the ratio of transconjugants to recipients.

### Competitive experiment of strains carrying type I or type II CRISPR-Cas systems

Each single colony was selected and placed in an LB tube with the corresponding antibiotic and cultured overnight at 37°C on a shaker. Bacteria from 1 mL culture was collected by centrifugation, then washed with LB twice and adjust to equal amount by OD_600_. Each strain was inoculated (1:100) into a new tube containing antibiotic-free LB, then grown in a shaker at 37°C for 12 h, the bacterial solution was serial diluted, and 50 μL of the bacterial solution was spread on the LB plate with appropriate antibiotics. The number of colonies on LB plate with or without antibiotics was counted. By comparing the percentages of different antibiotic resistant colonies, their growth competitiveness can be analyzed.

### Data availability

Raw CFU counts and conjugation frequency evaluation are available in the source data file. The sequences of plasmids used in this study were deposited on GenBank.

