## Supplemental Material for "Engineering probiotic *Escherichia coli* Nissle 1917 to block transfer of multiple antibiotic resistance genes by exploiting a type I CRISPR-Cas system"

**SUPPLEMENTAL MATERIALS AND METHODS**

**Figure S1**

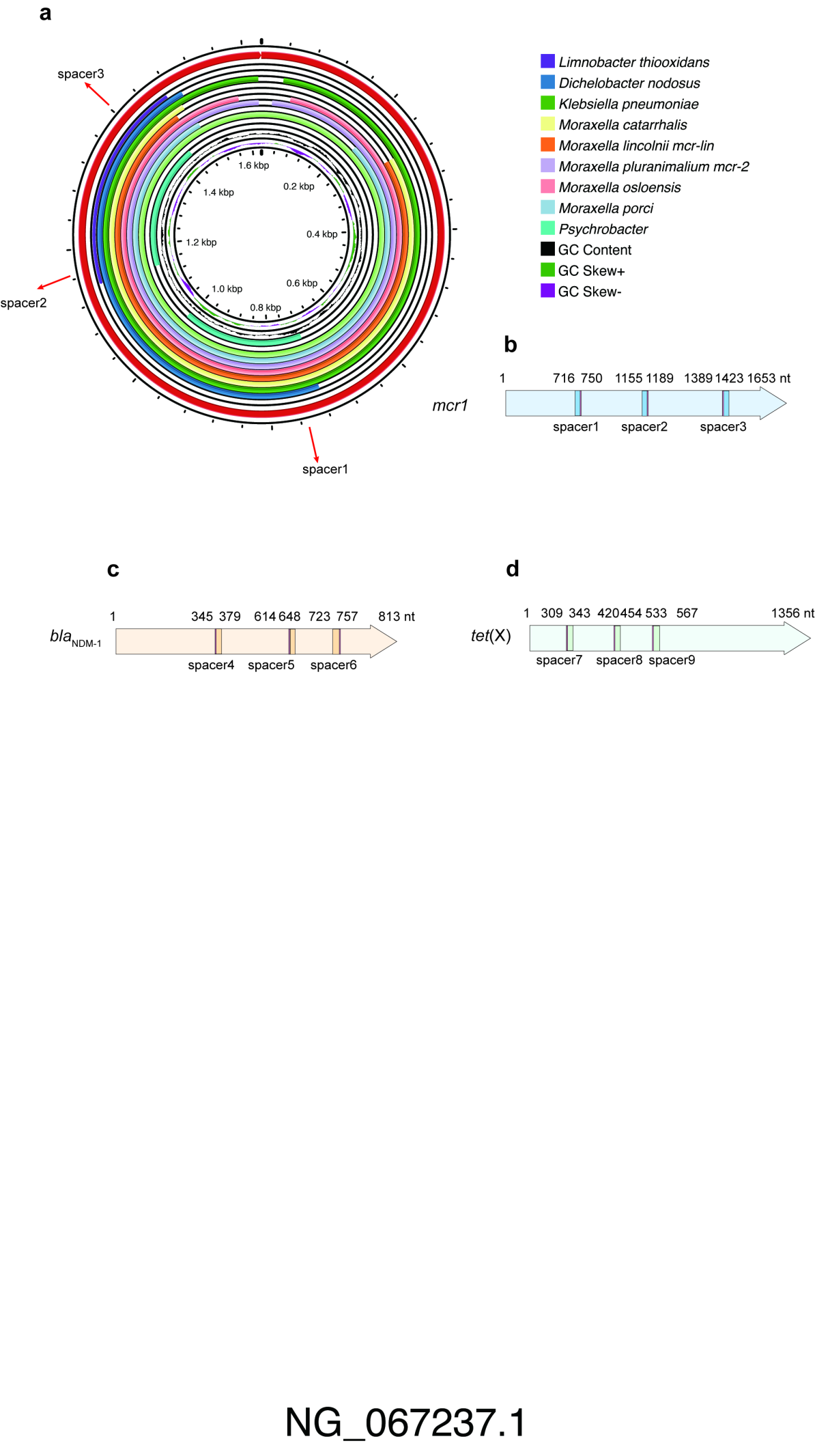

(**a**) Comparative analysis of *mcr* genes using *Escherichia coli mcr-*1 gene (NG_067237.1) as the reference (red circle). The Proksee (https://proksee.ca/) was used to visualize the conserved region to select spacers, which were illustrated with red arrows. The detailed information of sequences were listed in source data. (**b**-**d**)Schematic of the spacer targeting regions in *mcr-1*, *bla*_NDM-1_, *tet*(X). Genes are depicted as arrows in different colours and the matched spacers are shown as colour boxes with PAM (red box).

**Table 1 | List of strains used in the study.**

| **Strain** | **Description^A^** | **Source** |
| --- | --- | --- |
| WM3064 | Donor strain for conjugation: *thrB1004 pro thi rpsL hsdS lacZ*ΔM15 RP4-1360 Δ(*araBAD*)*567* Δ*dapA1341*::[*erm pir*(*wt*)] | ^1^ |
| *Escherichia coli* Nissle 1917 | Wild type of the probiotic E. coli Nissle 1917 | Forhigh Biotech Co., Ltd |
| DH5α | F^-^ø80(*lacZ*) ∆M15 ∆(*lacZYA*-*argF*) U169 *endA*1 *recA*1 *hsdR*17 (r_k_^-^, m_k_^+^ *sup*E44λ- *thi*-1 *gyrA*96 *relA*1 *phoA* | Tsingke Co., Ltd. |
| BW25113 | *lacI*^q^ *rrnB*_T14_ ∆ *lacZ*_WJ16_ *hsdR514* ∆*araBAD*_AH33_ ∆*rhaBAD*_LD78_ | ^2^ |
| ZJUTCBB0009 | BW25113 Δ*hns* | ^3^ |
| EC101 | BW25113Δ*hns* Δ*cas3* | This study |
| EC102 | BW25113Δ*hns* Δ*cas* | This study |
| E20-2 | Carrying *mcr-1* | This study |
| E119 | Carrying *bla*_NDM-1_ | This study |
| ZJ19PC | Carrying *tet*(X) | ^4^ |
| EcN-Cas3 | *cas3* induced by arabinose promoter was integrated into EcN genome | This study |
| EcN-Cas3-Cas | *cas3* and *cas* operon induced by arabinose promoter was integrated into EcN genome | This study |

**^A^ Gene designations are Cm coding for chloramphenicol resistance, Km for kanamycin resistance. FRT is the Flp recombinase target site. Resistance cassettes flanked by FRT-sites were deleted using temperature-sensitive plasmid pCP20.**

**Table 2 | List of plasmids used in the study.**

| **Plasmids** | **Description** | **Reference or source** |
| --- | --- | --- |
| pKD46 | Expressing Red recombinase, *repA101*(Ts) *oriR101*, Ap^R^ | ^5^ |
| pCP20 | Expressing FLP recombinase, *repA101*(Ts) pSC101 *ori*, Cm^R^, Ap^R^ | ^5^ |
| pKD3 | FRT Cm^R^ PS1 PS2 *ori*R6K, Ap^R^ | ^5^ |
| pKD4 | FRT Km^R^ PS1 PS2 *ori*R6K, Ap^R^ | ^5^ |
| pHG101 | Conjugation plasmid, Kan^R^ | A gift from Pro. Yin |
| pGLO | pUC 18/19 multiple cloning site, Amp^R^ | Lab reserve |
| pGLO-P_BAD_-*gfp* | pGLO derivative, *gfp* expressed from P_BAD_, Amp^R^ | Lab reserve |
| pGLO-J23100-*gfp* | pGLO derivative, GFP expressed by J23100, Amp^R^ | This study |
| pGLO-*leuO*-CR_MNT_ | pGLO derivative, carrying *leuO* expressed by P_BAD_ and a CRISPR array targeting to *mcr-1*, *bla*_NDM-1_, *tet*(X), Amp^R^ | This study |
| pGLO-CR_MNT_ | pGLO derivative carrying a CRISPR array targeting to *mcr-1*, *bla*_NDM-1_, *tet*(X), Amp^R^ | This study |
| T-*mcr-1* | pMD19-T-vector inserted with *mcr-1* gene, Amp^R^ | This study |
| T-*bla*_NDM-1_ | pMD19-T-vector inserted with *bla*_NDM-1_ gene, Amp^R^ | This study |
| T-*tet*(X) | pMD19-T-vector inserted with *tet*(X) gene, Amp^R^ | This study |
| pSU-P_BAD_-*leuO* | pSU19 derivative, carrying *leuO* expressed from P_BAD_, Cm^R^ | Lab reserve |
| pSU-Cas3 | pSU19 derivative, carrying *cas3* expressed from P_BAD_, Cm^R^ | This study |
| pSU-Cas | pSU19 derivative, carrying *cas* operon expressed from P_BAD_, Cm^R^ | This study |
| pDsRED | Non-Targeted plasmid expressing red fluorescent protein, pUC19 derivative, pMB1 replicon, Amp^R^ | Lab reserve |
| pCRI | pDsRED derivative carrying 4 spacers targeted by the type I-E CRISPR-Cas system of *E. coli*, Amp^R^ | Lab reserve |
| pGLO-Cas3-CR_MNT_ | pGLO derivative, Cas3 expressed by P_BAD_, CR_MNT_ expressed by J23100, Amp^R^ | This study |
| pHG101-*mcr-1* | pHG101 derivative carrying *mcr-1*, Kan^R^ | This study |
| pHG101-*bla*_NDM-1_ | pHG101 derivative carrying *bla*_NDM-1_, Kan^R^ | This study |
| pHG101-*tet*(X) | pHG101 derivative carrying *tet*(X), Kan^R^ | This study |
| pSU-*cas9* | pSU19 derivative, *cas9* expressed by P_BAD_, Cm^R^ | This study |

**Table 3 | List of oligonucleotides used in this study**

| **Project/task** | **No.** | **Primer** | **Sequence (5’→ 3’) ^A^** | **Description** |
| --- | --- | --- | --- | --- |
| Cloning | P1 | Replace P_BAD_ with J23100-F | TTGACGGCTAGCTCAGTCCTAGGTACAGTGCTAGCACCCGTT  TTTTTGGGCTAG | Constructing pGLO-J23100-*gfp* |
|  | P2 | Replace P_BAD_ with J23100-R | GCTAGCACTGTACCTAGGACTGAGCTAGCCGTCAAGGTCCC  GCTTTGTTACAG |  |
|  | P3 | Clone CR_MNT_ to pGLO-F | TCCTAGGTACAGTGCTAGCGTGGGTTGTTTTTATGGGA | Constructing pGLO-CR_MNT_ |
|  | P4 | Clone CR_MNT_ to pGLO-R | CTTCCCCCCGGTATCAACAGGGATCCACCTATGTCTGAACTC |  |
|  | P5 | Linearize pGLO -F | **GTTGCTGATAACCATCCT**TGCAAACCCTATGCTACTCC |  |
|  | P6 | Linearize pGLO -R | **ACATTTTCAGTGACCACGAC**GAATCCCTGCTTCGTCCA |  |
|  | P7 | Clone *leuO* to pGLO-F | **TTAAGAAGGAGATATACAT**ATGCCAGAGGTACAAAC | Constructing pGLO-*leuO*-CR_MNT_ |
|  | P8 | Clone *leuO* to pGLO-R | **GGTACCGAGCTCGAATTC**TTAGCGTTTGCAAATTGAGAC |  |
|  | P9 | Linearize pGLO-CR_MNT_ -F | ATGTATATCTCCTTCTTAAAGTTAAACAA |  |
|  | P10 | Linearize pGLO-CR_MNT_ -R | GAATTCGAGCTCGGTACCC |  |
|  | P11 | Clone *mcr-1* -F | atgATGCAGCATACTTCTGT | Constructing T-*mcr-1* |
|  | P12 | Clone *mcr-1* -R | tcaGCGGATGAATGCGG |  |
|  | P13 | Clone *bla*_NDM-1_ -F | atgGAATTGCCCAATATTATGCAC | Constructing T-*bla*_NDM-1_ |
|  | P14 | Clone *bla*_NDM-1_ -R | caGCGCAGCTTGTCG |  |
|  | P15 | Clone *tet*(X) -F | atgACTTTACTAAAACATAAAAAAATTACA | Constructing T-*tet*(X) |
|  | P16 | Clone *tet*(X) -R | ttaTAGATTCATTAGTTTTTGGAAAGAAAA |  |
|  | P17 | *cas3*-F | atgGAACCTTTTAAATATATATGCC | Constructing pSU-*cas3* |
|  | P18 | *cas3*-R | ttaTTTGGGATTTGCAGGGA |  |
|  | P19 | Linearize pSU-PBAD-LeuO -F | **ATATATTTAAAAGGTTCcat**ATGTATATCTCCTTCTTAAAGT  TAAACAAAATTATTTC |  |
|  | P20 | Linearize pSU-PBAD-LeuO -R | **TCCCTGCAAATCCCAAAtaa**GAATTCGAGCTCGGTACC |  |
|  | P21 | *cas-*F | atgAATTTGCTTATTGATAACTGGA | Constructing pSU-*cas* |
|  | P22 | *cas*-R | tcaCAGTGGAGCCAAAGATA |  |
|  | P23 | Linearize pSU-*cas3* -F | **TCCAGTTATCAATAAGCAAATTcat**ATGTATATCTCCTTCTTAAAGTTAAACAAAATTATTTC |  |
|  | P24 | Linearize pSU-*cas3* -R | **ATCTTTGGCTCCACTGtga**GAATTCGAGCTCGGTACC |  |
|  | P25 | *cas9-F* | **AGAAGGAGATATACAT**atgGATAAGAAATACTCAATAGGC | Constructing pSU-*cas9* |
|  | P26 | *cas9-R* | **CGGGTACCGAGCTCGAA**TTCtcaGTCACCTCCTAGCTG |  |
|  | P27 | Linearize pSU-*cas* -F | ATGTATATCTCCTTCTTAAAGTTAAACAAAATTA |  |
|  | P28 | Linearize pSU-*cas* -R | GAATTCGAGCTCGGTACC |  |
|  | P29 | Clone *mcr-1* to pHG-F | **TGACCGTGTGCTTCTCAA**atgATGCAGCATACTTCTGT | Constructing pHG101-*mcr-1* |
|  | P30 | Clone *mcr-1* to pHG-R | **ACCTGAAATTCAGTCGACC**tcaGCGGATGAATGCGG |  |
|  | P31 | Clone *bla*_NDM-1_ to pHG-F | **TGACCGTGTGCTTCTCAA**atgGAATTGCCCAATATTATGCAC | Constructing pHG101-*bla*_NDM-1_ |
|  | P32 | Clone *bla*_NDM-1_to pHG -R | **ACCTGAAATTCAGTCGACC**tcaGCGCAGCTTGTCG |  |
|  | P33 | Clone *tet*(X) to pHG-F | **TGACCGTGTGCTTCTCAA**atgATGACTTTACTAAAACATA | Constructing pHG101-*tet*(X) |
|  | P34 | Clone *tet*(X) to pHG-R | **ACCTGAAATTCAGTCGACC**ttaTAGATTCATTAGTTTTTGGAAAGA |  |
|  | P35 | Linearize pHG101 -F | CATTTGAGAAGCACACGGTC |  |
|  | P36 | Linearize pHG101 -R | CCTGAGGCCAGTTTGCTC |  |
|  | P37 | Clone P_BAD_-*cas3*-F | **ACTCCCGCCATTCAGAG**AAGAAACCAATTGTCCATATTGC | Constructing pGLO-*cas3*-CR_MNT_ |
|  | P38 | Clone P_BAD_-*cas3*-R | **AGGACTGAGCTAGCCGTCAA**AAGCTTGCATGCCTGCA |  |
|  | P39 | Linearize pGLO-CR_MNT_ -F | CTCTGAATGGCGGGAGT |  |
|  | P40 | Linearize pGLO-CR_MNT_ -R | TTGACGGCTAGCTCAGTC |  |
| Inactivation of chromosomal  Gene | P41 | CmFRT-F | ATTAGCCATGGTCCATATGAAT |  |
|  | P42 | CmFRT-R | CGATTGTGTAGGCTGGAG |  |
|  | P43 | *cas3* upstream-F | CTGCTCCATGAAAACTACCA | Generating donor for *cas3* inactivation |
|  | P44 | *cas3* upstream-R | **ATATGGACAATTGGTTTCTT**CAGCCGAAACATCAGTATTC |  |
|  | P45 | *cas3* downstream-F | TCATATGGACCATGGCTAATATGGCAAGCCTCATTCTC |  |
|  | P46 | *cas3* downstream-R | CACTCAGCTTCATAGAACGT |  |
|  | P47 | *cas* genes upstream-F | CGTATATTTTGGCGGCTCA | Generating donor for inactivating *cas* genes |
|  | P48 | *cas* genes upstream-R | ATATGGACAATTGGTTTCTTGTTGAGGAGACGATCTCATT |  |
|  | P49 | *cas* genes downstream-F | TCATATGGACCATGGCTAATAGCAGAGAAGAGGTCTTGAC |  |
|  | P50 | *cas* genes downstream-R | GGTTGAGGGGCATAATCAC |  |
| Intergration of gene into chromosome | P51 | Intergrate P_BAD_-*cas3*-F | AAGAAACCAATTGTCCATATTGC |  |
|  | P52 | Intergrate P_BAD_-*cas3*-R | **CTCCAGCCTACACAATCGT**TATTTGGGATTTGCAGGGATG |  |
|  | P53 | Intergrate P_BAD_-*cas*-F | AAGAAACCAATTGTCCATATTGC |  |
|  | P54 | Intergrate P_BAD_-*cas*-R | **CTCCAGCCTACACAATCGT**GCGAAACCCGTGTACCA |  |
| Genotyping | P53 | Test-cas3 intergration-F | CCGGCTTAACCGATAAGT | Checking *cas3* integration in EcN |
|  | P54 | Test-cas3 intergration-R | CGTAAACAGGCGATGTATAC |  |
|  | P55 | Test-cas intergration-F | CAGCAACGAACACGTGAA | Checking *cas* genes integration in EcN |
|  | P56 | Test-cas intergration-R | ACGGTAGATTTTATCCGGATG |  |
|  | P57 | Test-hns-F | AGTAAGTATTGGCTGCCAAT | Checking *hns* inactivating in BW25113 |
|  | P58 | Test-hns-R | CCACTTGTCGATAAGCCATT |  |
|  | P59 | Test-cas3-F | GCCCCAGGCGATATTTCTAT | Checking *cas3* inactivating in BW25113 |
|  | P60 | Test-cas3-R | GCGTACAGGGATCCAGTTAT |  |
|  | P61 | Test-cas-F | GGAAGGGAAACAGAATCTGG | Checking inactivation of *cas* genes in BW25113 |
|  | P62 | Test-cas-R | AATGGGAGCTGGGAGTTCTA |  |

**^A^ Homologou region in primers for assembly of DNA sequences by one-step cloning**

**SUPPLEMENTAL METHODS**

**Plasmid construction**

1. Constructing pGLO-J23100-*gfp*

The arabinose promoter located upstream of *gfp* was replaced with the constitutive promoter J23100 by T-PCR using primers P1/P2.

2. Constructing T-*mcr*-1, T*-bla*_NDM_-1, and T-*tet*(X), pHG101-*mcr*-1, pHG101-*bla*_NDM-1_, and pHG101-*tet*(X)

The *mcr*-1, *bla*_NDM_-1, and *tet*(X) genes were amplified with primer pairs P11/P12, P13/P14, P15/P16, respectively, followed by adding an A-tail (Takara Co., LTD, 6109) and inserting into pMD19-T vector by using a TA cloning kit (Takara Co., LTD, 6013).

The *mcr*-1, *bla*_NDM_-1, and *tet*(X) genes were amplified with primer pairs P29/P30, P31/P32, P33/P34, respectively. pHG101 was linearized with primers P35 / 36. pHG101-*mcr*-1, pHG101-*bla*_NDM-1_, and pHG101-*tet*(X), was constructed by recombining *mcr*-1, *bla*_NDM_-1, and *tet*(X) with linearized pHG101.

3. Constructing pSU-*cas3*, pSU-*cas*, pSU-Cas9 plasmid

The pSU-*cas3* plasmid was constructed by recombining the *cas3* gene (amplified from BW25113 genome with primer pair P17/P18) and the linearized pSU-PBAD-LeuO (using primers P19/P20). The pSU-*cas* plasmid was constructed by recombining the *cas* genes (*cse1*, *cse2*, *cas7*, *cas5*, *cas6,* amplified from BW25113 genome with primer pair P21/P22) and the linearized pSU-*cas3* (using primers P23/P24). The pSU-*cas9* plasmid was constructed by recombining the *cas9* gene (amplified with primer pair P25/P26) and the linearized pSU-*cas* (using primers P27/P28).

4. Constructing pGLO-CR_MNT_, pGLO-*leuO*-CR_MNT_ , pGLO-*cas3*-CR_MNT_

CR_MNT_ was synthesized as a CRISPR array containing 9 spacers selected from the sequences of *mcr-1*, *bla*_NDM_-1, and *tet*(X) (each 3 spacers). DNA synthesis were performed by Tsingke Biotech Co., LTD.

To construct pGLO-CR_MNT_, CR_MNT_ was amplified with primers P3/P4, and the pGLO vector was linearized with primers P5/P6. The linearized vector was recombined with the CR_MNT_ fragments by using the One Step Cloning Kit (Vazyme Co., LTD). pGLO-*leuO*-CR_MNT_ was constructed by recombining *leuO* gene (amplified from pSU-P_BAD_-*leuO* with primers P7/P8) and the linearized pGLO-CR_MNT_ (using primers P9/P10). pGLO-*cas3*-CR_MNT_ was constructed by recombining P_BAD_-*cas3* (amplified from pSU-*cas3* with primers P37/P38) and the linearized pGLO-CR_MNT_ (using primers P39/P40).
